# DeepSignal: detecting DNA methylation state from Nanopore sequencing reads using deep-learning

**DOI:** 10.1101/385849

**Authors:** Peng Ni, Neng Huang, Feng Luo, Jianxin Wang

**Affiliations:** School of Information Science and Engineering, Central South University, Changsha, 410083, China.; School of Computing, Clemson University, Clemson, SC, 29634, USA

## Abstract

The Oxford Nanopore sequencing enables to directly detect methylation sites in DNA from reads without extra laboratory techniques. In this study, we develop DeepSignal, a deep learning method to detect DNA methylated sites from Nanopore sequencing reads. DeepSignal construct features from both raw electrical signals and signal sequences in Nanopore reads. Testing on Nanopore reads of pUC19, E. coli and human, we show that DeepSignal can achieve both higher read level and genome level accuracy on detecting 6mA and 5mC methylation comparing to previous HMM based methods. Moreover, DeepSignal achieves similar performance cross different methylation bases and different methylation motifs. Furthermore, DeepSignal can detect 5mC and 6mA methylation states of genome sites with above 90% genome level accuracy under just 5X coverage using controlled methylation data.

## Introduction

DNA methylation, as a crucial form of epigenetic marks, plays important roles in a number of key biological processes [1, 2]. N6-methyladenine (6mA) and 5-methylcytosine (5mC) are the two most prevalent and well-studied base methylations. 5mC usually plays a role in embryonic development [3], atherosclerosis [4], aging and diseases [5]; and 6mA is important in transcriptional regulation [6], cancer development [7] and neurodevelopment [8].

Recently, the single molecule sequencing technologies, such as PacBio single molecule real time (SMRT) sequencing and Nanopore sequencing, are demonstrated to be able to detect the DNA methylation marks directly. Both technologies distinguish modified bases from standard nucleotide bases based on their distinctive signals. For PacBio SMRT sequencing, base modification would affect DNA polymerase kinetics, and then can be detected through different interpulse duration [9]. However, the accuracy of SMRT sequencing for detecting DNA methylation is heavily affected by the sequence coverage [10, 11]. Meanwhile, it also has been found that electrical signals in Nanopore sequencing are sensitive to epigenetic changes in the nucleotides [12, 13, 14]. Several studies have demonstrated that Nanopore sequencing can be used to detect DNA methylation.

Both statistics based and model based methods have been developed to identify base methylation from Nanopore sequencing reads. Simpson *et al*. [15] proposed a hidden Markov model (HMM) based approach to detect 5mC in CpG from events of Nanopore reads. The HMM model trained from E. coli data can detect CpG methylation in human Nanopore reads with 87% accuracy at read level. Due to the limitation in training data, the HMM method of Simpson et al. was not able to identify non-CpG methylation or a mixture of methylated and unmethylated CpGs. Rand *et al*. [16] also proposed an HMM-HDP based tool, called signalAlign, to classify 5mC at the inner cytosine of CCWGG motifs and 6mA at GATC motifs from events of Nanopore reads. singalAlign achieved 86%-95% accuracies at genome level for the Nanopore R9 data of pUC19 and E. coli. McIntyre *et al*. [17] used four kinds of classifiers (neural network, random forest, naive Bayes and logistic regression) to detect 6mA in mouse, E. coli and Lambda phage data. They achieved 84% accuracy at read level and 94% accuracy at genome level under 15X or higher coverage. Stoiber *et al*. [18] adopted Mann-Whitney U-test [19] in MoD-seq to detect controlled methylation in E coli. They achieved 0.839-0.896 genome level accuracies for seven different 5mC and 6mA motifs. Liu *et al*. [20] used Kolmogorov-Smirnov test [21] in their NanoMod tool to identify modified bases. Testing on E. coli methylation data, NanoMod have 70% precision at 50% recall at genome level. Both statistics based method can detect base modification in the absence of training dataset. However, to identify the modified bases in a native DNA, both methods require a matched amplified DNA as a control sample. Furthermore, statistics based methods achieved less accuracy comparing to model based methods.

Here, we present a deep learning method, called DeepSignal, to detect DNA methylated bases from Nanopore sequencing reads (Figure 1). DeepSignal employs two modules to construct features from raw electrical signals of Nanopore reads (Figure 1). First, since the methylation of a base does not only affect the signals of itself, but also affect the signal of its neighbors [20], the *signal* feature module in DeepSignal use convolutional neural network (CNN) to construct features directly from raw electrical signals around methylated base. Second, the methyltransferase need to bind to certain motif around the methylated site. To catch the sequence information around the methylated site, the sequence feature module in DeepSignal use bidirectional recurrent neural network (BRNN) to construct features from sequences of signal information. Then, features built from two modules are concatenated and fed into a fully connected neural network to predict the methylation states. We evaluate DeepSignal using both 5mC and 6mA data and the results show that DeepSignal achieves higher accuracy at both read level and genome level and requires less coverage of reads than previous HMM based methods do.

**Figure 1.**
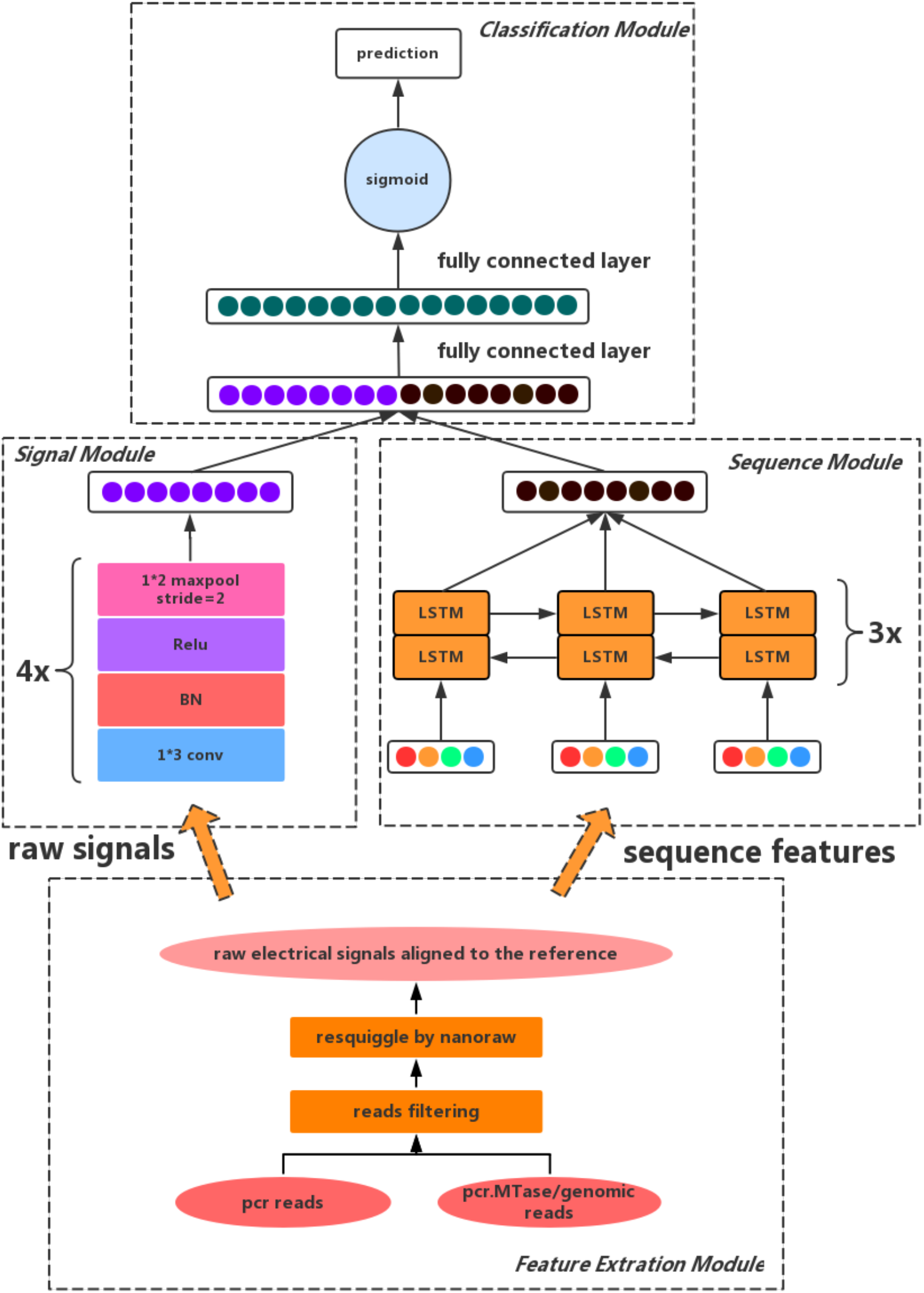
DeepSignal model for detecting methylation states

## Results

### Methylation signal in Nanopore reads

Previous studies [18] already showed that it is possible to distinguish the raw electrical signals of methylated site from those of unmethylated site using statistical tests. Figure 2 shows the boxplots of signals in methylated and unmethylated CpG and CCWGG of E. coli. The difference between raw signals of targeted base C is not significant between methylated and unmethylated reads. Meanwhile, the raw signal distribution of bases around targeted base C showed significant difference between methylated and unmethylated reads. Although statistics based method can identify signal difference of bases around targeted base, they ignored the relationship between bases. In this study, we design a deep learning method that try to catch both the signal and sequence information of bases around targeted base.

**Figure 2.**
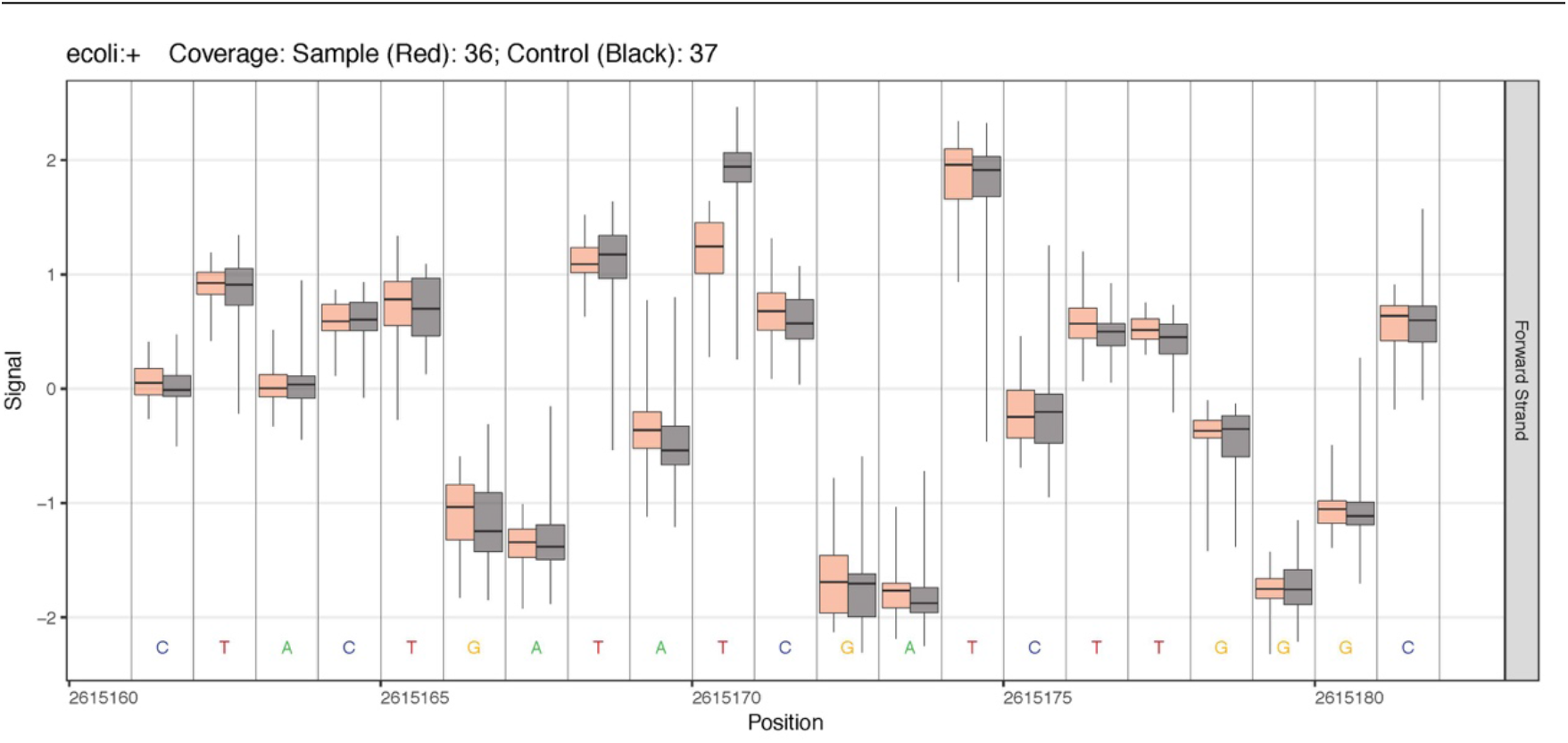
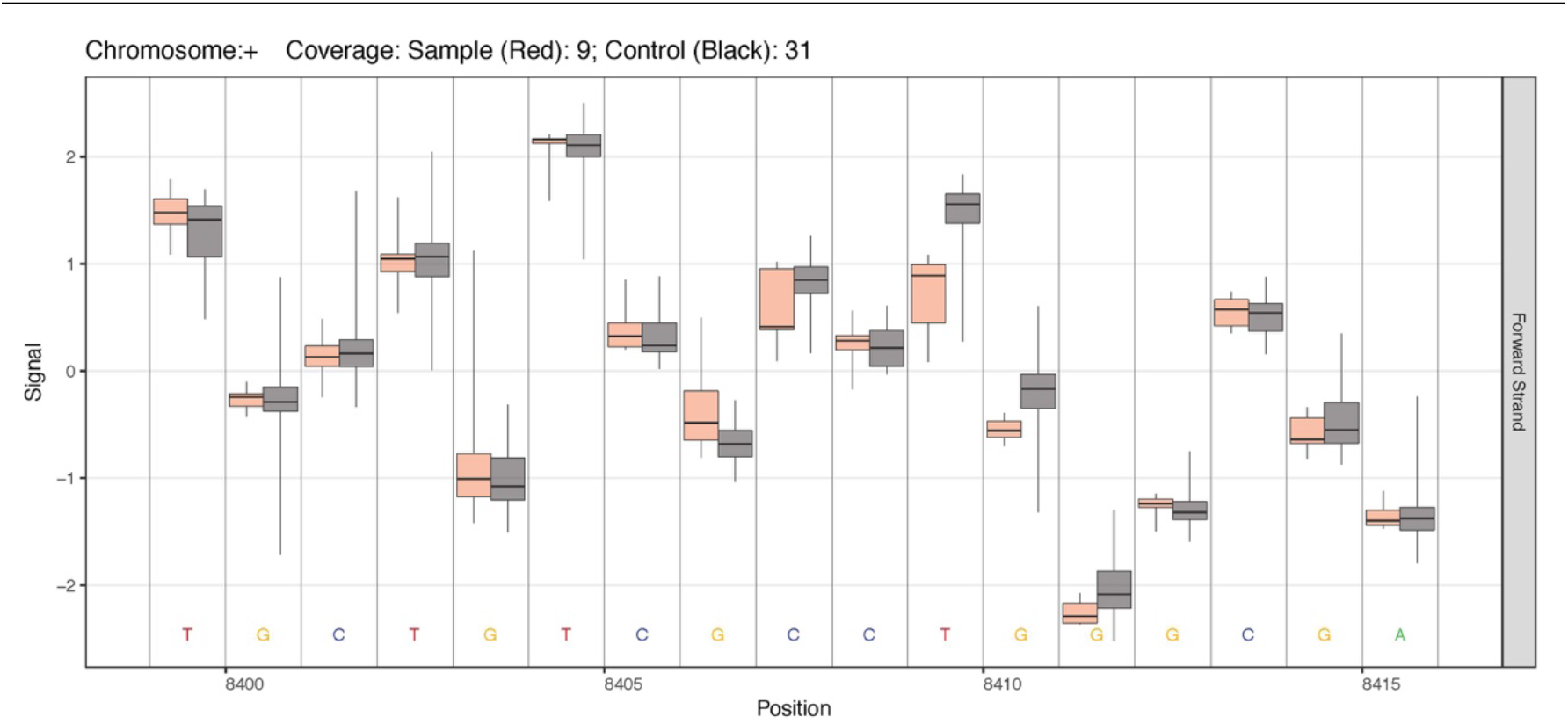
Boxplots of signal distribution of each base in methylated and unmethylated CpG and CCWGG of E. coli. A) CpG, B) CCWGG. The reads are from E. coli R9 2D data

### Evaluation of DeepSignal on G*A*TC methylation data of pUC19 plasmid

We first test DeepSignal using GATC methylation (6mA) data of pUC19 plasmid [16], which contains Nanopore reads of *dcm* methyltransferase treated pUC19 vector DNA and its PCR-amplified control. After extracting signal using nanoraw [18], we use 40% of reads for training both DeepSignal and signalAlign. Then, we test both models by randomly selecting 40 methylated and 40 unmethylated reads from the remaining 60% reads. We repeat our testing experiments 100 times. For each targeted site in reference genome, the methylation state is summarized from the predicted probabilities of all reads aligned to this site. When the model trained from template strand data is used, DeepSignal achieve 100% accuracy while signalAlign just get accuracy of 0.922 (Table 1). When both models trained from template and complement strand data are used, DeepSignal still achieve 100% accuracy while the accuracy of signalAlign is reduced to 0.906 (Table 1). Specially, DeepSignal achieves much higher specificities than siganlAlign does.

**Table 1.** Performance comparison of DeepSignal and signalAlign on classifying 6mA in GATC motifs of pUC19 DNA. Values are average and standard deviation of 100 replicated tests.

| strand | method | accuracy | Sensitivity | specificity | Precision |
| --- | --- | --- | --- | --- | --- |
| template | DeepSignal | 1.000(+/-0.000) | 1.000(+/-0.000) | 1.000(+/-0.000) | 1.000(+/-0.000) |
|  | signalAlign | 0.922(+/-0.024) | 0.962(+/-0.027) | 0.882(+/-0.042) | 0.892(+/-0.034) |
| both | DeepSignal | 1.000(+/-0.000) | 1.000(+/-0.000) | 1.000(+/-0.000) | 1.000(+/-0.000) |
|  | signalAlign | 0.906(+/-0.030) | 0.986(+/-0.020) | 0.826(+/-0.053) | 0.852(+/-0.039) |

### Evaluation of DeepSignal on C*C*WGG methylation data of pUCl9 plasmid

We also test DeepSignal using the CCWGG methylation (5mC) data of pUC19 plasmid, which contains Nanopore reads of *dam* methyltransferase treated pUC19 vector DNA and its PCR-amplified control. We also use 40% of reads for training both DeepSignal and signalAlign. Then, we randomly select 40 methylated and 40 unmethylated reads from the remaining 60% reads for testing. The performances are also evaluated on per targeted site of reference genome. When the model trained from template strand data is used, DeepSignal achieves accuracy of 0.996 while signalAlign just gets accuracy of 0.980 (Table 2). When both models trained from template and complement strand data are used, DeepSignal still achieve 100% accuracy while the accuracy of aignalAlign is 0.965. Similar to the performance on GATC methylation data, DeepSignal achieve similar sensitivities, but higher specificities.

**Table 2.** Performance comparison of DeepSignal and signalAlign on classifying 5mC in CCWGG motifs of pUC19 DNA. Values are average and standard deviation of 100 replicated tests

| strand | method | accuracy | sensitivity | specificity | Precision |
| --- | --- | --- | --- | --- | --- |
| template | DeepSignal | 0.996(+/-0.014) | 0.992(+/-0.027) | 1.000(+/-0.000) | 1.000(+/-0.000) |
|  | signalAlign | 0.980(+/-0.028) | 0.995(+/-0.026) | 0.965(+/-0.050) | 0.968(+/-0.045) |
| both | DeepSignal | 1.000(+/-0.005) | 0.999(+/-0.010) | 1.000(+/-0.000) | 1.000(+/-0.000) |
|  | signalAlign | 0.965(+/-0.038) | 1.000(+/-0.000) | 0.929(+/-0.075) | 0.938(+/-0.063) |

### Evaluation of DeepSignal on cross organism prediction of CpG methylation states

To further evaluate the performance of DeepSignal, we test it using CpG methylation (5mC) data of E. coli and human [15], which is previously studied by nanopolish. Both data include Nanopore reads of PCR-amplified DNA that was either untreated or treated with CpG methyltransferase M.SssI. We use nanoraw [18] to extract signals from Nanopore reads. In order to compare with nanopolish, we only select CpG sites have signals in both template and complement strands. There are 49378592 CpG sites for E. coli genome and 3120754 CpG sites for human genome. We train DeepSignal and nanopolish using 49378592 CpG sites of E. coli genome and test them on 3120754 CpG sites of human genome. Both DeepSignal and nanopolish learn two models from template and complement data separately. During testing, we predict the targeted site on template and complement separately using different models. As in nanopolish, we only output the final prediction on template strand, which is summarized by predicted probabilities of methylation states on both strand. The performances of both tools are summarized per-read level. As shown in Table 3, DeepSignal achieve 0.949 accuracy, while nanopolish only get an accuracy of 0.894. While DeepSignal has a relatively higher specificity comparing to nanopolish (0.970 vs. 0.947), DeepSignal has much higher sensitivity comparing to nanopolish (0.930 vs. 0.844). This result shows that DeepSignal is more powerful than nanopolish in classifying CpG sites.

**Table 3.** Performance comparison of DeepSignal and nanopolish on classifying CpGs in human

| method | accuracy | sensitivity | specificity | precision |
| --- | --- | --- | --- | --- |
| DeepSignal | 0.949 | 0.930 | 0.970 | 0.970 |
| nanopolish | 0.894 | 0.844 | 0.947 | 0.944 |

### Evaluation of the data coverage effect on DeepSignal

The coverage of reads usually affects the methylation state prediction. We evaluate DeepSignal with three experiments of testing on sampled sub-set of reads. First, for human CpG data, we count the coverage of targeted sites and evaluate the accuracies of methylation states with different coverage separately. Table 4 shows that DeepSignal achieves higher accuracy than nanoplish does for all 5 coverage levels. Even with 1X coverage level, DeepSignal still has 0.922 accuracy, while nanopolish has 0.875. The performance of both DeepSignal and nanopolish get improved when the coverage is increased from 1X to 3X. The performances of both tools decrease at 4X and 5X coverage may due to dramatically reduced targeted sites.

**Table 4.** Accuracies of DeepSignal and nanopolish for classifying CpGs in human R9 2D data in reference level under different coverages

| coverage threshold | 1 | 2 | 3 | 4 | 5 |
| --- | --- | --- | --- | --- | --- |
| (number of CpG sites) | (2895001) | (137330) | (4681) | (282) | (65) |
| DeepSignal | 0.922 | 0.968 | 0.970 | 0.929 | 0.938 |
| nanopolish | 0.875 | 0.932 | 0.942 | 0.894 | 0.815 |

For CCWGG and GATC methylation data of pUC19 plasmid, we evaluate the coverage effect on DeepSignal by sampling different number of reads. As shown in table 5, when the model trained from template strand data is used for predict GATC methylation in pUC19, DeepSignal can achieve 0.951 accuracy with only 5 sample reads, which signalAlign need 30 sample reads to have above 0.9 accuracy. When both models trained from template and complement strand data are used for predict GATC methylation in pUC19, DeepSignal can achieve 0.911 accuracy with only 2 sampled reads, which signalAlign need even 40 sample reads to have above 0.906 accuracy.

Table 6 shows that DeepSignal can achieve 0.920 accuracy with only 5 sampled reads when the model trained from template strand data is used for predict CCWGG methylation in pUC19, which signalAlign need 10 sample reads to have 0.920 accuracy. When both models trained from template and complement strand data are used for predict CCWGG methylation in pUC19, DeepSignal can achieve 0.952 accuracy with only 5 sampled reads, which signalAlign also need 10 sample reads to have above 0.9 accuracy.

**Table 5.** Accuracies of DeepSignal and signalAlign for classifying 6mA in GATC motifs of pUC19 DNA under different number of sampled reads.

| strand | method | Number of sampled reads |  |  |  |  |  |  |
| --- | --- | --- | --- | --- | --- | --- | --- | --- |
|  |  | 1 | 2 | 5 | 10 | 20 | 30 | 40 |
| template | DeepSignal | 0.866<br>(+/-0.082) | 0.912<br>(+/-0.062) | 0.951<br>(+/-0.030) | 0.983<br>(+/-0.018) | 0.997<br>(+/-0.007) | 1.000<br>(+/-0.003) | 1.000<br>(+/-0.000) |
|  | signalAlign | 0.738<br>(+/-0.084) | 0.750<br>(+/-0.078) | 0.793<br>(+/-0.047) | 0.843<br>(+/-0.042) | 0.887<br>(+/-0.037) | 0.911<br>(+/-0.031) | 0.922<br>(+/-0.024) |
| both | DeepSignal | 0.838<br>(+/-0.061) | 0.911<br>(+/-0.046) | 0.966<br>(+/-0.028) | 0.992<br>(+/-0.012) | 0.999<br>(+/-0.005) | 1.000<br>(+/-0.002) | 1.000<br>(+/-0.000) |
|  | signalAlign | 0.686<br>(+/-0.064) | 0.729<br>(+/-0.060) | 0.810<br>(+/-0.051) | 0.850<br>(+/-0.044) | 0.882<br>(+/-0.034) | 0.898<br>(+/-0.029) | 0.906<br>(+/-0.030) |

**Table 6.** Accuracies of DeepSignal and signalAlign for classifying 5mC in CCWGG motifs of pUC19
DNA under different number of sampled reads.

| strand | method | Number of sampled reads |  |  |  |  |  |  |
| --- | --- | --- | --- | --- | --- | --- | --- | --- |
|  |  | 1 | 2 | 5 | 10 | 20 | 30 | 40 |
| template | DeepSignal | 0.834<br>(+/-0.125) | 0.863<br>(+/-0.103) | 0.920<br>(+/-0.060) | 0.969<br>(+/-0.040) | 0.985<br>(+/-0.024) | 0.995<br>(+/-0.017) | 0.996<br>(+/-0.014) |
|  | signalAlign | 0.795<br>(+/-0.134) | 0.832<br>(+/-0.104) | 0.862<br>(+/-0.085) | 0.920<br>(+/-0.065) | 0.956<br>(+/-0.038) | 0.972<br>(+/-0.034) | 0.980<br>(+/-0.028) |
| both | DeepSignal | 0.842<br>(+/-0.093) | 0.888<br>(+/-0.081) | 0.952<br>(+/-0.054) | 0.981<br>(+/-0.034) | 0.994<br>(+/-0.017) | 0.995<br>(+/-0.016) | 1.000<br>(+/-0.005) |
|  | signalAlign | 0.747<br>(+/-0.102) | 0.797<br>(+/-0.091) | 0.864<br>(+/-0.073) | 0.916<br>(+/-0.054) | 0.940<br>(+/-0.047) | 0.955<br>(+/-0.040) | 0.965<br>(+/-0.038) |

## Methods

### Nanopore sequencing data

#### CpG methylation (5mC) data

The Nanopore reads of methylated and unmethylated CpGs of E. coli K12 ER2925 and human NA12878 previously used by Simpson *et al*. are downloaded from the European Nucleotide Archive under accession PRJEB13021. Only the R9 2D data with raw signals are used in our study.

#### C<u>C</u>WGG methylation (5mC) and G*A*TC methylation (6mA) Data

The Nanopore reads of methylated and unmethlylated pUC19 plasmid DNA (both 5mC and 6mA data) and the E. coli K12 MG1655 (5mC data) previously used by signalAlign [16] are downloaded from NCBI under accession SRP098631.

### DeepSignal Model

DeepSignal is made up of four modules (Figure 1). The first module extracts raw signals from Nanopore reads and prepares the input data for feature construction modules. We have designed two modules to extract different features. The *signal* feature module that constructs features from raw electrical signals around modified base using convolutional neural network (CNN) and the *sequence* feature module that constructs features from the sequence of signal information around modified base using bidirectional recurrent neural network (BRNN). Finally, a classification module of neural network takes features built from two modules and predicts the methylation states.

### Signal extraction from Nanopore reads

To extract signals from Nanopore reads, we use the re-squiggle module of nanoraw [18] to map the raw electrical signal values to contiguous bases in the genome reference. The nanoraw corrects the indel errors in Nanopore base call and re-annotates raw signal to match the genomic bases. After the re-squiggle, we extract corresponding raw signals of each matched base in the genome reference. The genome references of E. coli and human used for alignment are E.coli K12 MG1655 and GRCh38.p5 respectively. The pUC19 plasmid reference sequence is downloaded from NCBI Nucleotide.

For each candidate of methylated site (a C in CpG or CCWGG, an A in GATC), we extract raw signal of 17 bp nucleotide sequence centering around it. Then, for each base, we calculate the mean, standard deviation, and length of its signal values. Therefore, for each sample, we have four 17-length vectors as input for following *sequence* module: the 17-mer nucleotide, the mean signal value, the standard deviation of signal values and the length of signal values. Meanwhile, we select 120 raw signal values from the middle of the 17-mer’s signal values as input to *signal* feature module.

### *Sequence* feature module

In *sequence* feature module, we train a bidirectional recurrent neural network (BRNN) to construct features from four 17-length signal information vectors. Each BRNN [1] include a forward RNN and backward RNN to catch both past and future context. A RNN scan the sequence of data and encodes the sequential information into hidden state vector h. We use the long short-term memory (LSTM) RNN in our BRNN. Let *x*_1_, *x*_2_,…, *x_t_* are a sequence of signal values (the 17-mer nucleotide, the mean signal value, the standard deviation of signal values and the length of signal values). A LSTM RNN will recursively calculate the hidden layer *h* as follows:

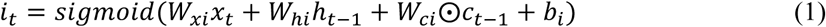

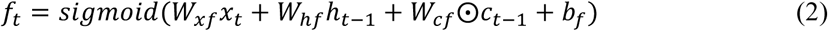

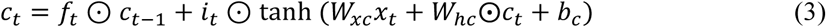

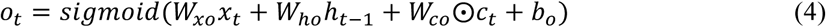

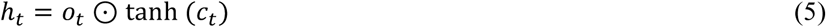

where *W* and *b* are weights and biases in the model. *x* is the input vector; *i* is the activation vector of input gate; *f* is the activation vector of forget gate; *c* is the cell state vector; *o* is the activation vector of output gate; and *h* is the output vector of the LSTM hidden unit. Current output *h_t_* of LSTM hidden unit depends on the input *x_t_*, previous state *h*_*t*−1_, and previous information stored in cell. Then, the output of forward and backward LSTM RNN is combined:

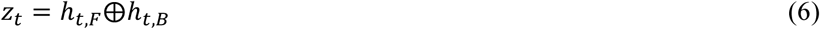

Our *sequence* feature module has three forward and three backward layers. For each sample, the *sequence* feature module generates a representation vector *z*, which will be inputted to classification module.

### *Signal* feature module

In *signal* feature module, we train a deep convolutional neural network (CNN) from 120 raw signal values of methylated site and its close neighbor sites. Our deep CNN stacks four building blocks. Each block include convolution, batch normalization, activation and pooling operations. First, the input vector *x* is transformed by 1d convolution operation:

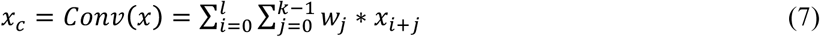

here *w* is the weight vector of the convolution kernel. Then, we apply batch normalization, which can keep the mean and variance fixed without saturation. The batch normalization over a batch *B* is:

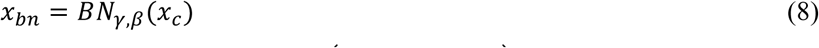

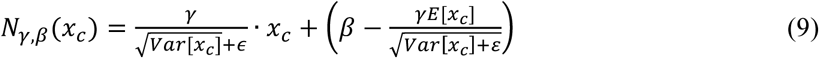

where *γ,β,ε* are the parameters to be learned. Batch normalization can improve the performance and stability of neural networks.

After batch normalization, the activation operation we used is *ReLu*, which sets negative value to zero:

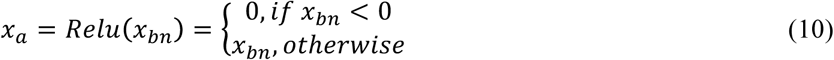

Finally, a pooling operation is used to summarize the activations of adjacent *k* neurons by taking their maximum value:

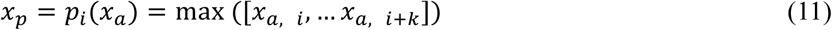

we sequentially select *k* neurons without overlapping for pooling.

### Classification Module

The classification module takes the output vectors from sequence and signal feature and concatenate them as the input. The high-level reasoning in classification module is done by a fully connected neural network with two hidden layers; each followed by a dropout layer to prevent overfitting. The output layer uses a sigmoid activation function to predict the methylation probability *ŷ* ∊ [0.1] of the target site:

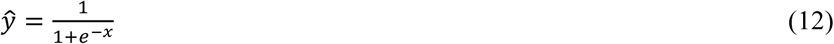

### Model training

Model parameters are learnt on the training set by minimize the cross-entropy loss function L as follows:

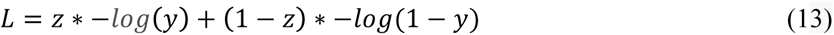

where *z* is the true label distribution of training data. We use a batch size of 512 and an initial learning rate 0.001. The learning rate is adapted by Adam [4] and decayed by a factor of 0.96 after each epoch. We keep the node with probability value of 0.5 at each dropout layer. The whole module in implemented in Python using tensorflow.

### Evaluation of DeepSignal

Since the template sequence and the complement sequence of Nanopore 2D reads are quite different in read quality and signal patterns [15], we train two models using template strand and complement strand separately.

#### Evaluation of DeepSignal using CpG methylation data

Same as Simpson *et al*., we use the E. coli data to train DeepSignal and test the model on human data. We select only reads with high mapping quality to reference genome (quality score greater than 20) to extract signal values [15]. However, even with high mapping quality reads, nanoraw still cannot align signals for some of reads. As a result, for human genome, we only have selected 3120754 CpG sites having signals in both template and complement strands. For E. coli genome, there are 49378592 CpG sites that have signals in both template and complement strands.

#### Evaluation of DeepSignal using GATC methylation data

We select the reads of which the 2D sequences cover more than 2600 bp of the pUC19 DNA reference. After the alignment using BWA, 12844 reads of gDNA and 18892 reads of pcrDNA are kept. Then, 40% of the reads in pUC19 data are used to train the DeepSignal model. Then, we randomly select 40 methylated and 40 unmethylated reads from the remaining 60% for testing and repeat 100 times [16]. We use the same reads to train and test DeepSignal and siganlAlign. The performances of models are evaluated in reference level as described in [16].

#### Evaluation of DeepSignal using CCWGG methylation data

For CCWGG methylation data of pUC19, we use the same procedure of above GATC methylation classification experiments.

#### Performance Evaluation of DeepSignal

We evaluate DeepSignal at per-read level and at genome site level. For each targeted base in a read, DeepSignal outputs two probabilities *P*+ and *P*_-_, which represent the probabilities of methylated and unmethylated state, respectively. If *P*_+_ > *P*_-_, the methylation state of targeted base is predicted as methylated, otherwise is predicted as unmethylated. Based on the predicted results, we calculate accuracy, sensitivity, specificity and precision of DeepSignal at per-read level as following:

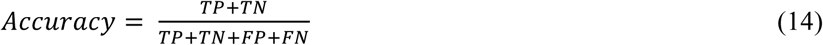

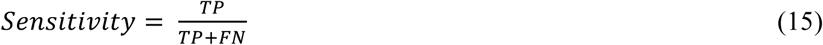

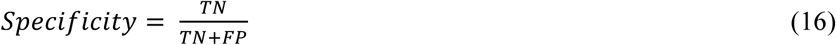

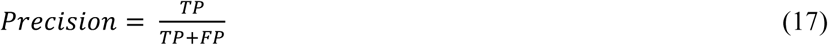

where *TP* is true positives, *TN* is true negatives, *FP* is false positives, *FN* is false negatives. To evaluate DeepSignal at genome level, we group the predictions of targeted bases in reads based on their aligned positions in reference genome. Then, we calculate the methylated probability *P*+ ’ and the unmethylated probability *P*_-_ ’ of each tested site in reference as following:

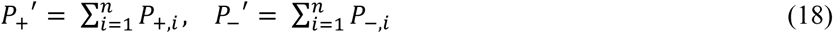

where *n* is the number of targeted bases in reads which are aligned to the site in reference genome, *P*_+, *i*_ and *P*_-,*i*_ are the methylated probability and the unmethylated probability of the targeted base in ith read. Then, we calculate the performance of DeepSignal at reference genome level using equations (14) - (17). We evaluate nanopolish and signalAlign using the same methods.

## Conclusion

In this study, we develop DeepSignal, a deep learning method to detect DNA methylation states from Nanopore sequencing reads. Testing DeepSignal on Nanopore reads of pUC19, E. coli and human, we show it can achieve higher accuracy on detect both 6mA and 5mC sites comparing to previous HMM based methods. DeepSignal is able to detect 5mC in both CpGs and CCWGG motifs. Furthermore, DeepSignal achieve similar performance on different methylation bases and different methylation motifs, while other methods, like signalAlign, has higher performance on 5mC methylation site than on 6mA methylation site.

Due to still relative high cost of Nanopore sequencing, it is important to identify methylation sites with low coverage of reads. We demonstrate that DeepSignal can detect 5mC and 6mA methylation site at genome level with above 90% accuracy under 5X coverage using controlled methylation data. DeepSignal requires much lower coverage than those required by HMM and statistics based methods.

## Acknowledgement

We are grateful to Miten Jain for providing their methylation data.

## References

[1] Schübeler,Dirk. “Function and information content of DNA methylation.” Nature 517.7534 (2015): 321.

[2] Bergman, Yehudit, and Howard Cedar. “DNA methylation dynamics in health and disease.” Nature structural & molecular biology 20.3 (2013): 274.

[3] Smith, Zachary D., and Alexander Meissner. “DNA methylation: roles in mammalian development.” Nature Reviews Genetics 14.3 (2013): 204.

[4] Lund, Gertrud, et al. “DNA methylation polymorphisms precede any histological sign of atherosclerosis in mice lacking apolipoprotein E.” Journal of Biological Chemistry 279.28 (2004): 29147–29154.

[5] Gonzalo, Susana. “Epigenetic alterations in aging.” Journal of applied physiology 109.2 (2010): 586–597.

[6] Yue, Yanan, Jianzhao Liu, and Chuan He. “RNA N6-methyladenosine methylation in post-transcriptional gene expression regulation.” Genes & development 29.13 (2015): 1343–1355.

[7] Xiao, Chuan-Le, et al. “N6-Methyladenine DNA Modification in the Human Genome.” Molecular Cell 71 (2018): 306-318 e7.

[8] Yao, Bing et al., Active N6-Methyladenine Demethylation by DMAD Regulates Gene Expression by Coordinating with Poly-comb Protein in Neurons, Molecular Cell (2018)

[9] Flusberg, Benjamin A., et al. “Direct detection of DNA methylation during single-molecule, real-time sequencing.” Nature methods 7.6 (2010): 461.

[10] Davis, Brigid M., Michael C. Chao, and Matthew K. Waldor. “Entering the era of bacterial epigenomics with single molecule real time DNA sequencing.” Current opinion in microbiology 16.2 (2013): 192–198.

[11] Zhu, Shijia, et al. “Mapping and characterizing N6-methyladenine in eukaryotic genomes using single-molecule real-time sequencing.” Genome research (2018).

[12] Schatz, Michael C. “Nanopore sequencing meets epigenetics.” Nature methods 14.4 (2017): 347.

[13] Laszlo, Andrew H., et al. “Detection and mapping of 5-methylcytosine and 5-hydroxymethylcytosine with nanopore MspA.” Proceedings of the National Academy of Sciences 110.47 (2013): 18904–18909.

[14] Schreiber, Jacob, et al. “Error rates for nanopore discrimination among cytosine, methylcytosine, and hydroxymethylcytosine along individual DNA strands.” Proceedings of the National Academy of Sciences (2013): 201310615.

[15] Simpson, Jared T., et al. “Detecting DNA cytosine methylation using nanopore sequencing.” nature methods 14.4 (2017): 407–410.

[16] Rand, Arthur C., et al. “Mapping DNA methylation with high-throughput nanopore sequencing.” Nature methods 14.4 (2017): 411.

[17] McIntyre, Alexa BR, et al. “Nanopore detection of bacterial DNA base modifications.” bioRxiv (2017): 127100.

[18] Stoiber, Marcus H., et al. “De novo identification of DNA modifications enabled by genome-guided nanopore signal processing.” bioRxiv (2016): 094672.

[19] Mann, Henry B., and Donald R. Whitney. “On a test of whether one of two random variables is stochastically larger than the other.” The annals of mathematical statistics (1947): 50–60.

[20] Liu, Qian, et al. “NanoMod: a computational tool to detect DNA modifications using Nanopore long-read sequencing data.” bioRxiv (2018): 277178.

[21] Massey Jr, Frank J. “The Kolmogorov-Smirnov test for goodness of fit.” Journal of the American statistical Association 46.253 (1951): 68–78.

